# DNA local structure decreases mutation rates

**DOI:** 10.1101/302133

**Authors:** Chaorui Duan, Qing Huan, Xiaoshu Chen, Shaohuan Wu, Lucas B. Carey, Xionglei He, Wenfeng Qian

**Affiliations:** State Key Laboratory of Plant Genomics, Institute of Genetics and Developmental Biology, Chinese Academy of Sciences, Beijing 100101, China; Human Genome Research Institute and Department of Medical Genetics, Zhongshan School of Medicine, Sun Yat-sen University, Guangzhou 510080, China; State Key Laboratory of Biocontrol, School of Life Sciences, Sun Yat-sen University, Guangzhou 510275, China; Key Laboratory of Genetic Network Biology, Institute of Genetics and Developmental Biology, Chinese Academy of Sciences, Beijing 100101, China; Department of Experimental and Health Sciences, Universitat Pompeu Fabra, Barcelona 08003, Spain; University of Chinese Academy of Sciences, Beijing 100049, China

**Author notes:** These authors contributed equally to this work (Chaorui Duan, Qing Huan, Xiaoshu Chen). Correspondence to: Wenfeng Qian, Institute of Genetics and Developmental Biology, Chinese Academy of Sciences, Beijing 100101, China.

**Keywords:** DNA shape, mutation rate, mutational landscape

## Abstract

**Background:** Mutation rates vary across the genome. Whereas many *trans* factors that influence mutation rates have been identified, as have specific sequence motifs at the 1-7 bp scale, *cis* elements remain poorly characterized. The lack of understanding why different sequences have different mutation rates hampers our ability to identify positive selection in evolution and to identify driver mutations in tumorigenesis.

**Results:** Here we show, using a combination of synthetic genes and sequencing of thousands of isolated yeast colonies, that intrinsic DNA curvature is the major *cis* determinant of mutation rate. Mutation rate negatively correlates with DNA curvature within genes, and a 10% decrease in curvature results in a 70% increase in mutation rate. Consistently, both yeast cells and human tumors accumulate mutations in regions with small curvature. We further show that this effect is due to differences in the intrinsic mutation rate, likely due to differences in mutagen sensitivity, and not due to differences in the local activity of DNA repair.

**Conclusions:** Our study establishes a framework in understanding the *cis* properties of DNA sequence in modulating the local mutation rate and identifies a novel causal source of non-uniform mutation rates across the genome.

## BACKGROUND

Mutation is the ultimate source of genetic diversity. Therefore, the measurement of mutation rate and particularly, the identification of the *trans* factors and *cis* elements that influence mutation rate are a focus of intense interest in evolutionary biology. A large number of *trans* factors influencing mutation rate have been identified [1], such as chromatin remodelers, histone-modifying enzymes, and other DNA binding proteins [2-4]. In addition, replication timing [5-9] and transcription rate [10-14] also affect mutation rate.

*Cis* elements may play a more important role in determining the local mutation rate, yet remain poorly understood. Studies of *cis* elements that determine local mutation rate have been limited to the scale of a few neighboring nucleotides around a mutation site for the past few decades [15-18].

There is comprehensive *cis* information in the shape of DNA. Although the double-helix structure of DNA is usually described as a twisted ladder, the steps of the ladder are not rigidly aligned. The local shape of DNA is affected by the interactions of neighboring bases [19, 20]. For example, the size of the minor and major grooves varies depending on the local sequence. Such variation in DNA shape affects the ability of proteins to bind to DNA and the accessibility of each nucleotide [20, 21]. Through its effect on DNA-protein and/or DNA-solvent interactions, the shape of the double helix may influence the local mutation rate. However, the role of DNA shape in influencing local mutation rate has not been systematically studied. Here, we provided several lines of evidence that intrinsic DNA curvature affects the local mutation rate in a quantitative and predictable manner. Our study therefore expands our knowledge of *cis* elements that regulate mutation rate by integrating information regarding the physical shape of the double helix and develops a new framework to understand the evolution of local mutation rate.

## RESULTS AND DISCUSSION

### Characterization of the mutational landscape of *URA3*

To quantitatively determine how *cis* elements affect the local mutation rate we first characterized the mutational landscape of an endogenous gene, *URA3*, in *Saccharomyces cerevisiae. URA3* encodes an enzyme required for uracil synthesis and converts the nontoxic molecule 5-fluoroorotic acid (5-FOA) into the toxic 5-fluorouracil. Only cells bearing loss-of-function mutations in *URA3* can survive on 5-FOA plates, making *URA3* a model gene to study mutation rate [5, 22]. Here, we cultured wild-type yeast in synthetic complete (SC) media for 24 hours to allow mutations to accumulate and spread these cells onto a 5-FOA plate (**Fig. 1A**). We then sequenced *URA3* of each randomly picked visible colony and identified mutations. We performed 135 biological replicates in parallel and sequenced a total of ∼1,000 *URA3* variants from 135 plates (**Table S1**). Identical mutations (same type at the same position) identified on the same plate were counted only once because such mutations are most likely resulted from cell proliferation from a single mutation and not independent identical mutations.

**Fig. 1.**
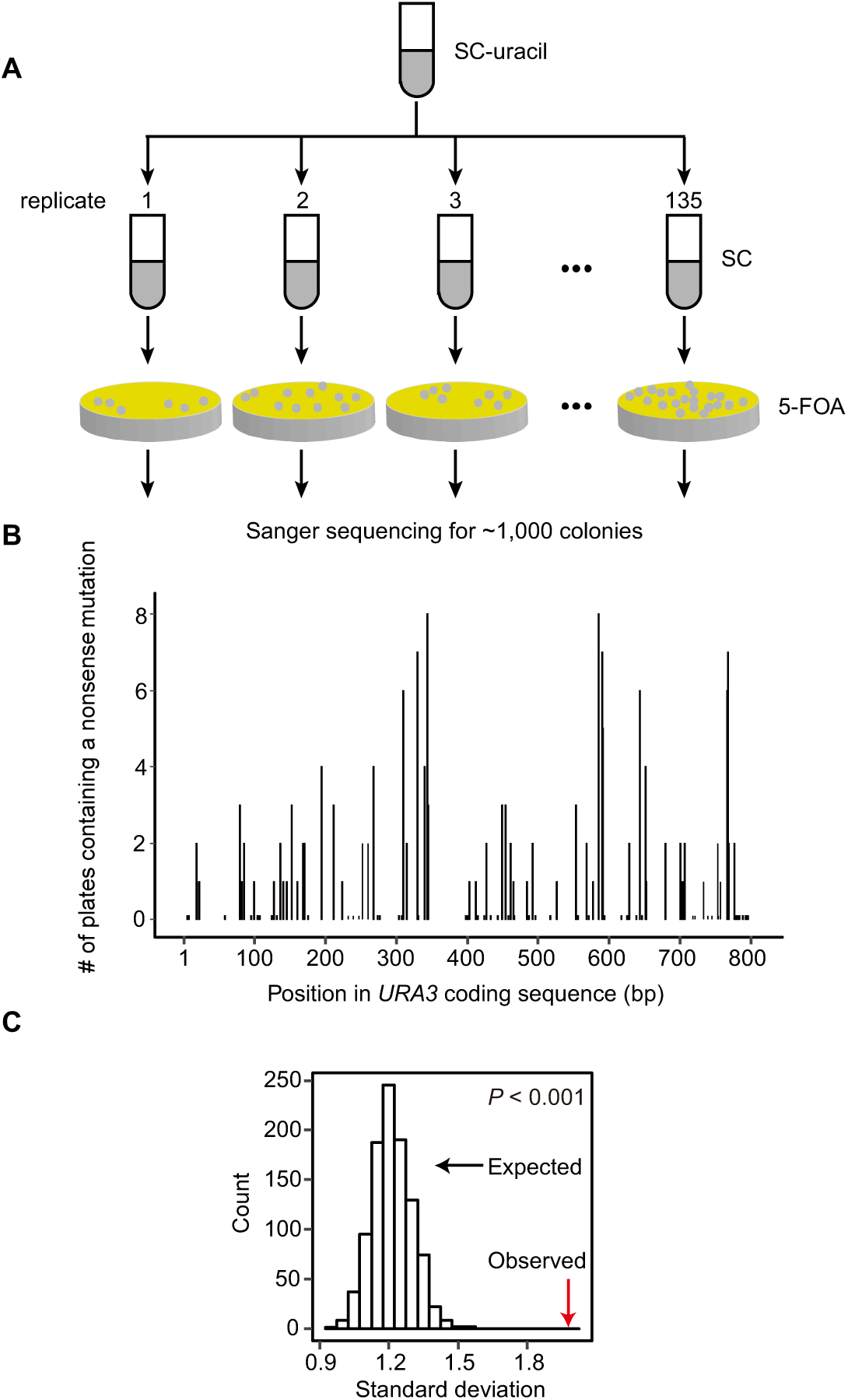
The mutational landscape of *URA3.* (A) A schematic description of the experimental design. Mutations were accumulated in SC liquid medium and *ura3* mutants were selected on 5-FOA plates. (B) Mutational landscape of all potential nonsense mutation sites, which were defined as sites where a point mutation can result in a stop codon. Each bar represents a potential nonsense mutation site. (C) The observed (red arrow) and expected (histogram showing 1,000 permutations) standard deviation of the numbers of nonsense mutations on all potential nonsense mutation sites of *URA3.*

To measure bias in mutation rate, we need to determine the number of observed mutations and to compare it with the number expected if the mutation rate was uniform. As the missense mutations that would permit growth on 5-FOA is unknown, we focused our analysis on nonsense mutants. There are 104 potential nonsense mutation sites in *URA3.* For each of them, we counted the number of 5-FOA plates where each nonsense mutation was observed (**Fig. 1B**). This number varied between 0 and 8 (**Fig. 1B**). To determine if this variation in frequency could be fully explained by the inherently stochastic nature of mutation, we randomly assigned each of the observed 154 nonsense mutations to a potential nonsense mutation site. We then calculated the standard deviation of the observed numbers of nonsense mutations on these sites and that in the permutation. The observed standard deviation was significantly greater than the random expectation (*P* < 0.001, **Fig. 1C**), suggesting the presence of *cis* elements that affect the local mutation rate.

A nonsense mutation may not always lead to a loss of function, especially when it occurs near the stop codon. This would also lead to a non-Poisson distribution of observed mutations. To exclude this confounding factor we repeated the permutation test using only the first two-thirds of the coding sequence. Again, the observed standard deviation was significantly greater than the random expectation (**Fig. S1A**). Similar results were also obtained when we performed the permutation test separately for the 54 nonsense transitions and the 100 nonsense transversions (**Fig. S1B-C**). Taken together, the variation in the frequency of nonsense mutations within *URA3* suggests the presence of *cis* elements that modulate local mutation rate.

### Mutations in *URA3* tend to occur in DNA regions with a smaller intrinsic DNA curvature

One possible explanation for the non-Poisson distribution of observed nonsense mutations is the difference in the mutation rate into a stop codon of each of the four bases. Nucleotides A and T had a lower mutation rate than G and C (**Fig. S2**), likely explained by the AT rich nature of the three stop codons. That is, G>A and C>T transitions often result in stop codons but A>G and T>C transitions do not. To explore the predictive power of the nucleotide at each position and to identify additional *cis* sequence features predictive of local mutation rates, we constructed a set of linear models that take into account various sequence features **(Table 1)**. Including the nucleotide at the potential nonsense site in the linear model decreases the Akaike information criterion (AIC) of the model, indicating an increase in the model’s ability to predict mutation rates (**Table 1**, model 1 and model 2). Surprisingly, including the +1 and –1 bases into the model did not further improve the predictive power, nor did including the heptanucleotide sequence context (**Table 1**, models 3 and 4).

**Table 1.** Models on predicting the mutation rate of a potential nonsense site in *URA3*.

|  | **Model** | **AIC** |
| --- | --- | --- |
| 1 | Null model | 484 |
| 2 | Mutation rate ∼ “0” * | 418 |
| 3 | Mutation rate ∼ “0” + “+1” + “-1” | 426 |
| 4 | Mutation rate ∼ “0” + “+1” + “+2” + “+3” + “-1” + “-2” + “-3” | 432 |
| 5 | Mutation rate ∼ curvature ** | 433 |
| 6 | Mutation rate ∼ “0” + curvature | 414 |

\* “0” represents the nucleotide at the potential nonsense site. “+1” and “-1” represent the upstream and the downstream nucleotide of the potential nonsense site, respectively.
\*\* The intrinsic DNA curvature in a region from 50 bp upstream to 50 bp downstream of the potential nonsense site.

To identify additional DNA sequence features predictive of local mutation rates we used a sliding window to divide the *URA3* gene into overlapping regions of *L* nucleotides (*L*= 10, 20 …, or 100 bp). We calculated the average mutation rate in each region as the total number of observed nonsense mutations in this region normalized by the number of potential nonsense mutation sites (**Fig. S3A**). For each region we then calculated 17 DNA properties such as GC content, thermodynamic characteristics, groove properties, and DNA shape features using well-established computational methods [19, 23] (**Fig. S3B**). Finally, for each window size, we calculated the correlation between mutation rate and each of the DNA properties.

Over a large range of window sizes, mutation rate was most strongly correlated with intrinsic DNA curvature, defined as the sequence-dependent deflection of DNA axis due to the interaction between neighboring base pairs [24] (e.g., for a window size *L* of 100 bp, *ρ* = −0.49, *P* = 2×10^−5^, Spearman’s correlation, **Fig. 2A-B**). Consistently, including intrinsic DNA curvature into the aforementioned linear model enhances its predictive power (**Table 1**, models 5 and 6). It is worth noting that tilt, the DNA property exhibiting the second strongest correlation with mutation rate, is a component of intrinsic DNA curvature [24].

**Fig.2.**
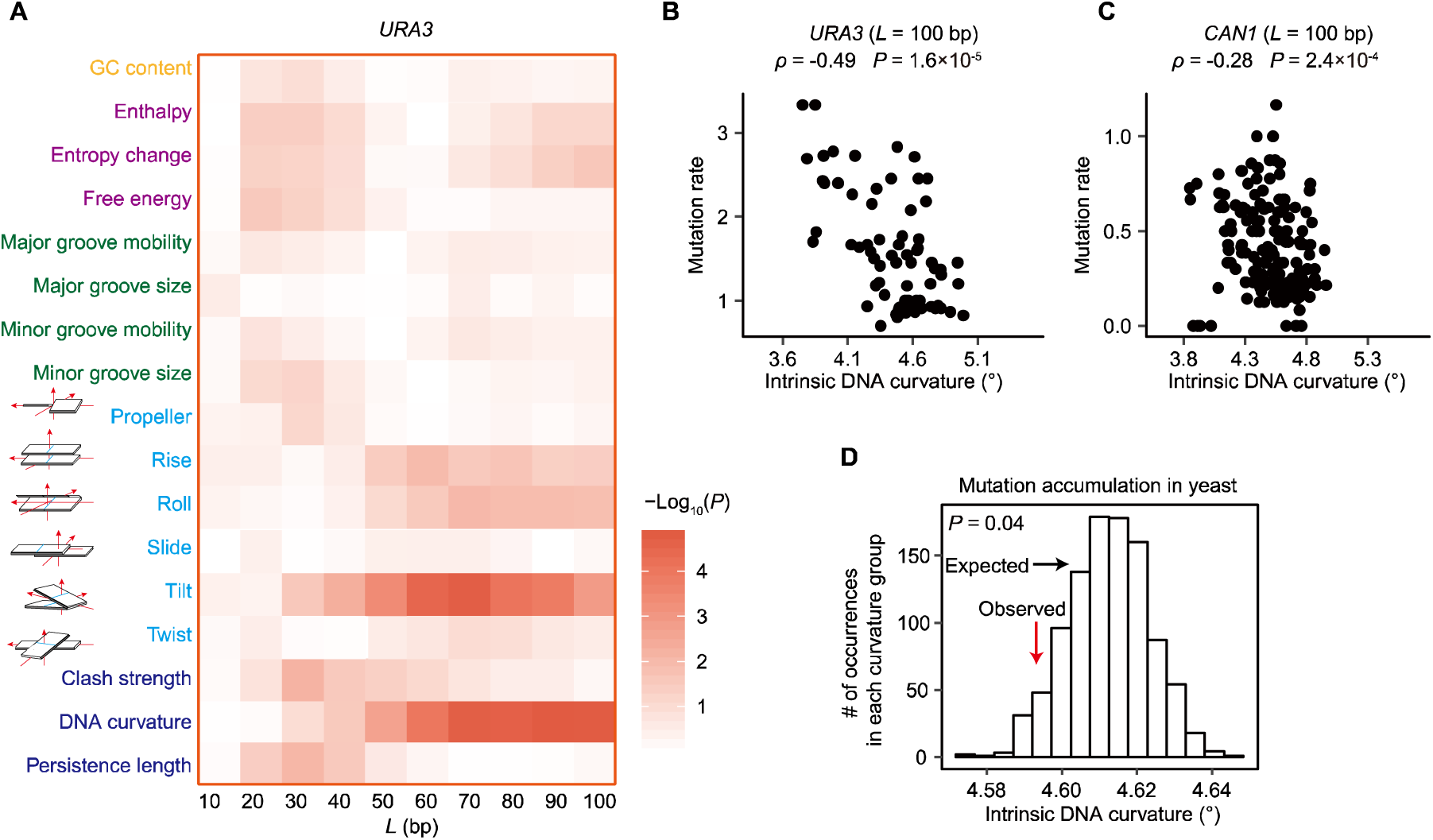
Intrinsic DNA curvature is negatively correlated with mutation rate. (A) Correlation between mutation rate and the value of each DNA feature in sliding windows (window length *L*). These features include GC content (orange), thermodynamic characteristics (purple), groove properties (green), intra- and inter-base pair DNA shape features (cyan), and integrated DNA shape features (blue). Intra- and inter-base pair DNA shape features are shown in cartoons, where a square represents a base and a rectangle represents a base pair. *P* values were calculated from the Spearman’s correlation. (B-C) Example scatter plots of *URA3* (B) and of *CAN1* (C). Each dot represents a region of length *L* (= 100 bp). (D) The average intrinsic DNA curvature of DNA regions surrounding the 882 observed mutation sites (red arrow) was significantly smaller than the random expectation (histogram showing 1,000 permutations) in the yeast genome. *P* value was calculated with a permutation test.

The correlation between mutation rate and DNA curvature was not confounded by GC content [17, 25] which in our data was not correlated with mutation rate (**Fig. 2A**). We previously showed that nucleosome binding suppresses spontaneous mutations [26]. To quantitatively determine the relationship between mutation rate, nucleosome occupancy, and DNA curvature, we performed high-throughput sequencing on nucleosome protected DNA fragments. Consistent with previous results, nucleosome occupancy was negatively correlated with the average mutation rate (*r_uRA3_*= –0.35, *P* = 0.002). Nevertheless, the correlation between DNA curvature and mutation rate persisted after controlling for nucleosome occupancy (partial *r_uRA3_* = –0.6, *P* = 1×10^−8^), suggesting that the relationship between mutation rate and DNA curvature is not due to differences in nucleosome occupancy.

As a form of experimental cross-validation to determine if our results from *URA3* are generalizable to other genes, we used an independently generated set of mutations in the gene *CAN1* [22], for which nonsense mutations were selected using the arginine analogue canavanine. Intrinsic DNA curvature is also predictive of mutation rate in *CAN1* (**Fig. 2C** and **Table S2**).

### Mutations in yeast cells and in human tumors accumulate in DNA regions with smaller intrinsic DNA curvature

To determine if DNA shape affects mutation rate at the genomic scale, we used a mutation accumulation assay in which spontaneous mutations accumulate at ∼100× the normal rate due to a mutation in a gene related to DNA mismatch repair, *MSH2* [27]. We retrieved all 882 mutations that were supported by an at least 20× coverage in the high-throughput sequencing data. We calculated the intrinsic DNA curvature of a region from 50 bp upstream to 50 bp downstream of each mutation. As a control we randomly chose 882 sites with identical 3-nucleotide contexts (the mutation site, +1, and –1 sites) from the rest of the genome. We performed this random sampling procedure 1,000 times. We found that the observed mutations were located in regions with a smaller intrinsic DNA curvature (*P* = 0.04, permutation test, **Fig. 2D**). It suggests that in the genome as a whole, regions with smaller intrinsic DNA curvature have higher mutation rates.

Mutations generate genetic variation among cells within multi-cellular individuals, and somatic mutations play a vital role in cancer development and progression. Mutations in tumors are distributed unevenly across the genome and within individual genes [2, 3, 9, 16, 28]. We therefore performed the same genome-scale analysis as in yeast using 10,429 cancer samples from 26 cancer types collected in The Cancer Genome Atlas (TCGA) database [29]. We calculated the average intrinsic curvature of the DNA regions from 50 bp upstream to 50 bp downstream of each identified SNP for each cancer type. As a control, we randomly chose the same number of DNA sites from the genome. Consistent with the results in yeast, mutations were significantly enriched in regions with a smaller intrinsic DNA curvature in all cancer types (*P* < 0.001, permutation test, **Fig. 3** and **Fig. S4**), suggesting that intrinsic DNA curvature reduces mutation rates in human tumor cells. The large number of mutations in tumor cells permitted a more robust test of the effect for nucleotide context. We found that DNA curvature negatively correlates with mutation rate when controlling for the trinucleotide (**Fig. S5**) or heptanucleotide context (**Fig. S6**). Taken together, DNA curvature is a robust predictor of non-uniform mutation rates in both yeast cells and human tumors.

**Fig. 3.**
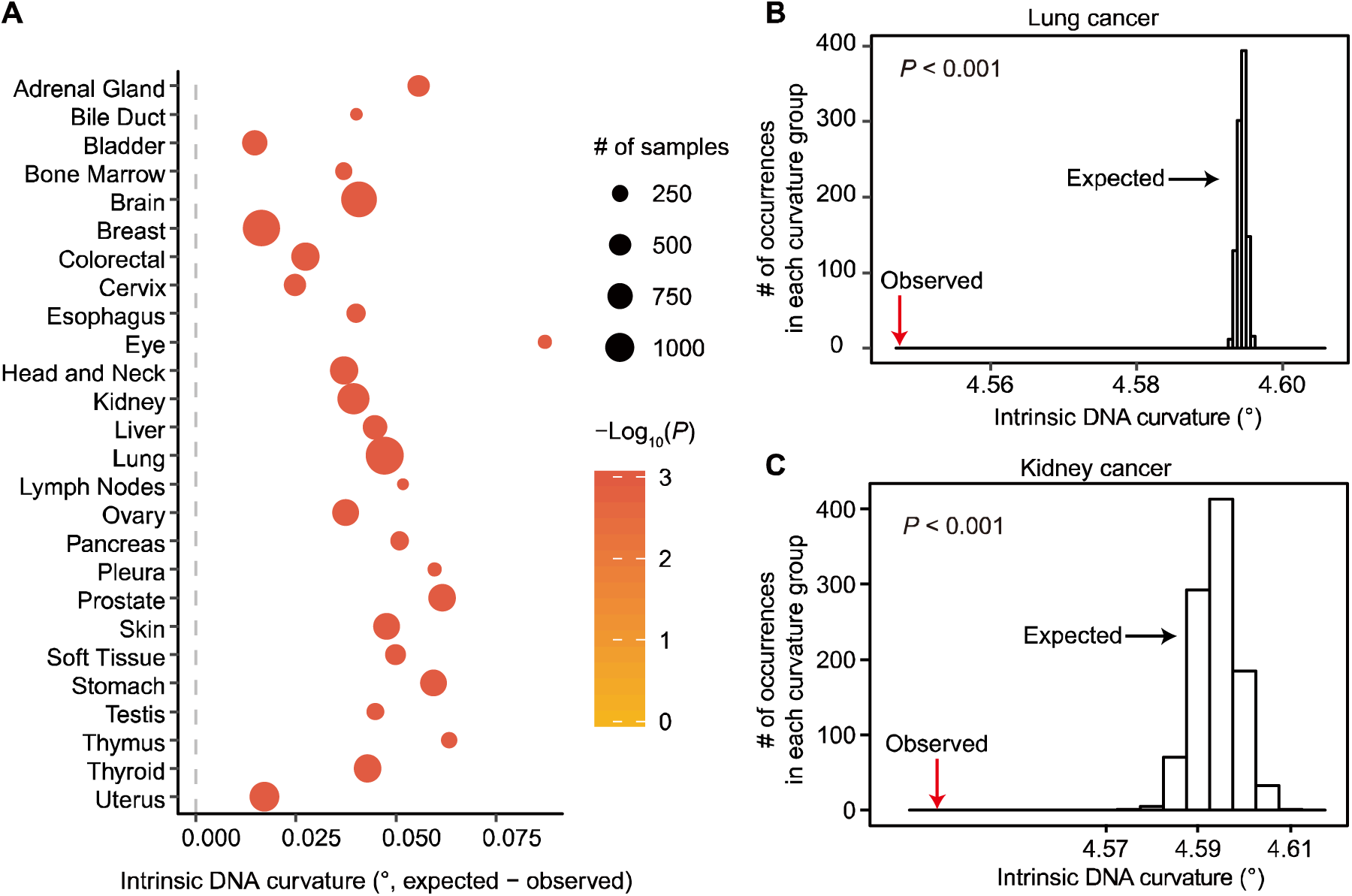
Mutations in human cancer samples are enriched in DNA regions with a smaller intrinsic DNA curvature. (A) Mutations are enriched in regions with a significantly smaller curvature in all 26 cancer types. Each dot represents a cancer type. *P* values were calculated based on the permutation test. *P* values were arbitrarily assigned to 0.001 when *P* < 0.001. (B-C) Examples in lung (B) and in kidney (C) showing that the average intrinsic DNA curvature of SNP-containing regions (red arrows) was significantly smaller than the random expectation (histogram showing 1,000 permutations).

### Genetic manipulation of DNA curvature affects mutation rate

To further examine the causal effect of intrinsic DNA curvature on mutation rate we designed four synonymous variants of *URA3* (**Table S3**), two with increased curvature and two with decreased curvature (**Fig. 4A**). We kept features that may influence local mutation rate such as GC content, codon usage, and predicted local mRNA structure largely unchanged (**Table S4**) [13, 17, 25]. The expression levels of *URA3* in these variants are also identical (**Fig. S7**).

**Fig. 4.**
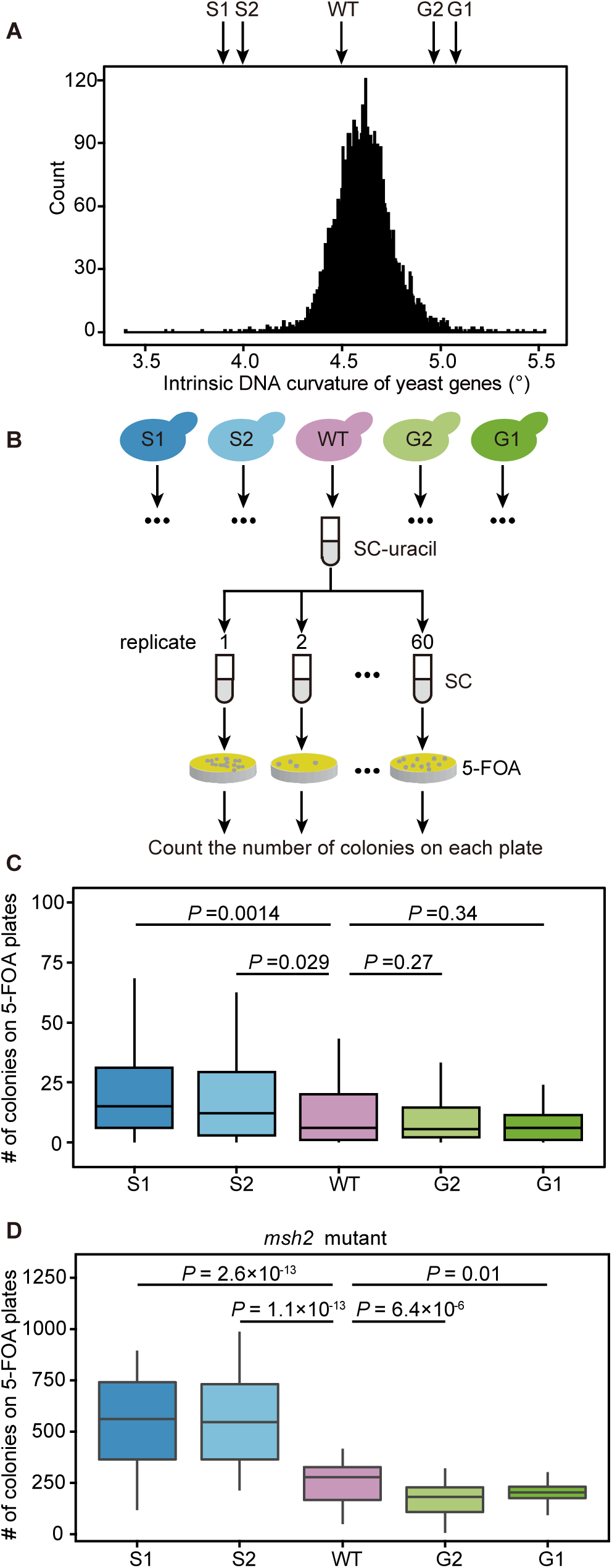
Changing the intrinsic DNA curvature in *URA3* leads to altered mutation rate. (A) The distribution of the average intrinsic DNA curvature of genes in the yeast genome. The intrinsic DNA curvatures of five synonymous variants of *URA3* are indicated by arrows. S1 and S2 (G1 and G2) are variants with a smaller (greater) intrinsic DNA curvature. (B) The schematic description of the experimental procedure for measuring the relative mutation rate of *URA3* variants. (C) Reduction of intrinsic DNA curvature leads to an increase in the mutation rate of *URA3.* Outliers are not shown. *P* values were calculated from the one-tailed Mann-Whitney *U* test. (D) Similar to (C), in a mismatch repair deficient *msh2* strain.

We used an electrophoretic mobility shift assay to confirm that the intrinsic DNA curvature was altered in these variants [30-32]. Variants with a greater predicted intrinsic DNA curvature [19, 23] migrated more slowly than those with a smaller curvature (**Fig. S8**), presumably due to the different friction force that they encountered in the process of migration.

To determine if genetic manipulation of curvature alters mutation rate we cultured cells with each of the five *URA3* variants in SC media to allow mutations to accumulate, spread cells onto 5-FOA plates, and counted the number of colonies on each plate (**Fig. 4B**). We calculated the mutation rate of each variant from the fraction of plates without mutants [33] and found that variants with a 10% smaller intrinsic DNA curvature had a 70% higher mutation rate (**Fig. 4C**). It suggests that experimental decreasing DNA curvature increases mutation rate.

### Intrinsic DNA curvature alters the mutation rate, not mismatch repair efficacy

There are two non-mutually exclusive mechanisms by which intrinsic DNA curvature can modulate the net mutation rate [9]. First, intrinsic DNA curvature may reduce the supply of mutations. Second, intrinsically curved DNA may facilitate the recruitment of mismatch repair-related proteins, which can increase the DNA repair efficacy [3, 9]. To determine if intrinsic DNA curvature reduces the supply of mutations or affects repair efficiency, we knocked out *MSH2* and repeated the mutation accumulation experiment (**Fig. 4B**). In the absence of Msh2, the effect of DNA curvature on mutation rate is even larger; a 10% decrease in curvature results in a 100% increase in mutation rate (**Fig. 4D**).

This observation suggests that the altered net mutation rate by DNA curvature is due to differences in the supply of mutations and not to differences in DNA repair efficacy.

### DNA curvature reduces mutagen sensitivity in cancer cells

DNA curvature may reduce the mutation rate by making the DNA sequence less accessible to potential mutagens [26] or by affecting the fidelity of DNA polymerase itself, though this is unlikely, as DNA polymerase acts on single stranded DNA. To distinguish these two mechanisms we divided the SNPs in cancer cells into six categories based on mutation types and asked if the rate of mutation types that are sensitive to mutagens is more affected by DNA curvature. C>T transitions mainly result from the hydrolytic deamination on methylated cytosine [15, 34]. The rate of C>T transition reduced by 40% in DNA regions with greater curvature (**Fig. 5A**). In contrast, this reduction in mutation rate was not observed for other mutation types (**Fig. 5A**). Furthermore, C>A transversions in lung cancer cells are mainly caused by polycyclic aromatic hydrocarbons in tobacco smoke [35-37]. C>A mutations are more affected by DNA curvature in lung cancer than they are in other types of cancer (**Fig. 5B**). Both biased distributions of C>T and C>A mutations suggest that curvature protects DNA from mutagens. Given the well-established role of DNA curvature in regulating protein-DNA interactions [20, 21], it is possible that DNA curvature promotes protein binding that makes DNA less accessible to mutagens.

**Fig. 5.**
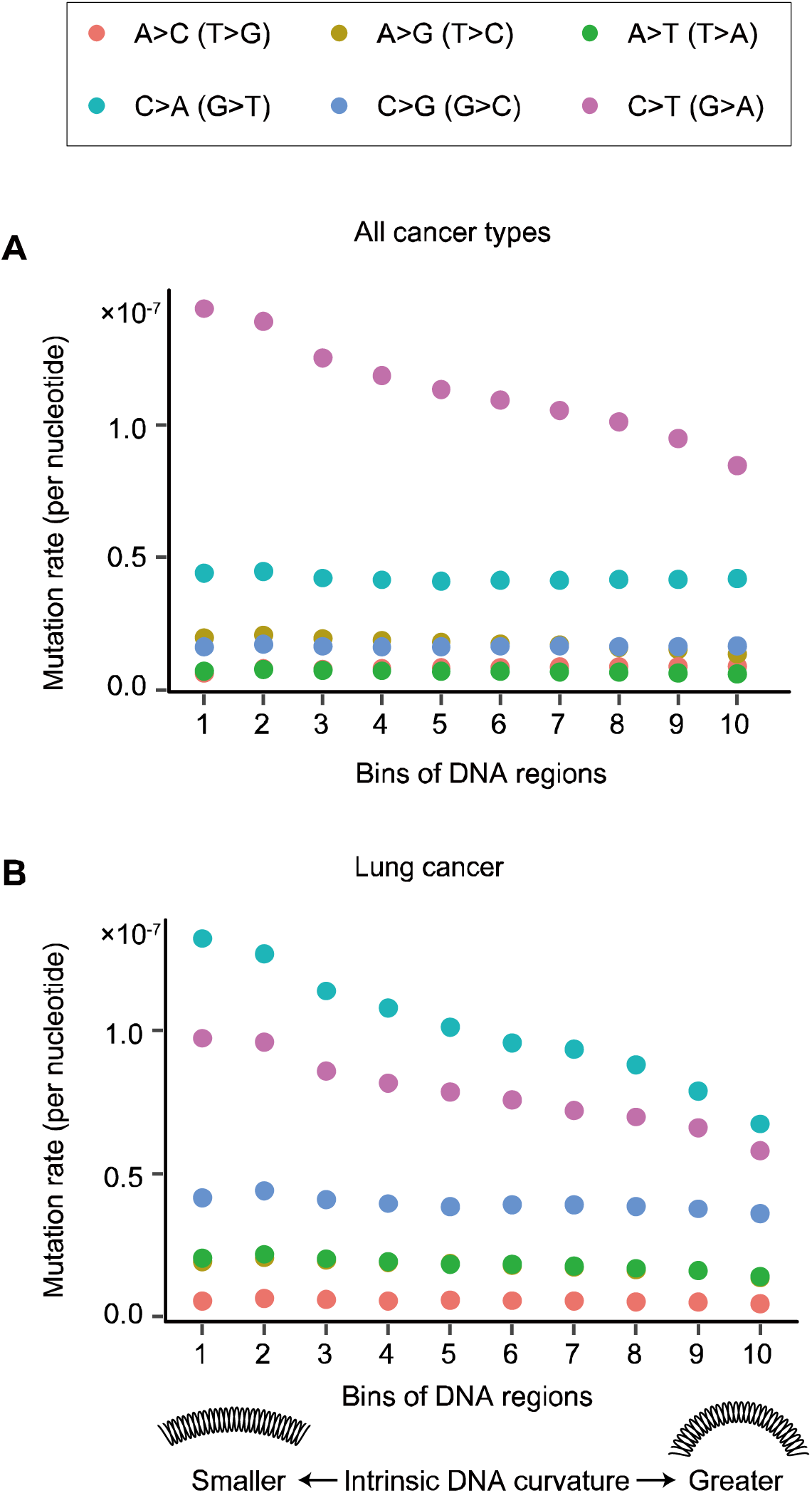
DNA curvature suppresses mutations that are induced by mutagens. (A) The mutation rate of each of the six mutation types in cancer cells. Mutation rate was defined as the number of SNPs per cancer sample per nucleotide. *x* axis shows ten equally sized bins of DNA regions in the human genome sorted by intrinsic DNA curvature. (B) Mutation rates in lung cancer (including lung adenocarcinoma and lung squamous cell carcinoma).

### Implications in evolutionary genomics

Understanding the variation in mutation rate is central to numerous questions in evolutionary genetics. Particularly, modeling the variability in mutation rate among sites of a genome is of key importance in studies of molecular evolution because it provides a null model that can be rejected when natural selection occurs. Sequence-intrinsic *cis* elements are more computationally tractable than *trans* factors in modeling mutation rate in molecular evolution studies, because with *cis* elements the expected mutation rate can be predicted directly from the surrounding sequences of a site [16]. For example, the evolutionary rates of genes have been extensively studied, and particularly, comparisons between those of essential and nonessential genes have been made [38-42]. Previous studies focused on the difference in the strength of negative selection and neglected the potential difference in mutation rate, presumably because the latter was hard to model. In this study, we discovered that a key DNA shape feature, intrinsic DNA curvature, modulated local mutation rate. Interestingly, we observed that essential genes exhibit a greater DNA curvature in both yeast (**Fig. S9**) and humans (**Fig. S10**), suggesting that they have a lower mutation rate. This observation urges the need of considering the difference in mutation rate when compare evolutionary rate among genes.

Furthermore, the high-density fitness landscapes of random mutations on a gene have been extensively characterized in previous studies [43, 44], aiming to understand the trajectory of biological evolution. However, evolutionary trajectories are determined by natural selection acting on mutations. Inherent biases in the generation of the random mutations must therefore be taken into account. Our study on mutational landscape complements these previous studies on fitness landscapes and will significantly contribute to the ultimate understanding of evolutionary trajectories [45].

## CONCLUSIONS

We found that the shape of the DNA double helix plays a major role in determining the local mutation rate. In particular, we identified a key feature, intrinsic DNA curvature, that determines the local mutation rate in both yeast and cancer cells. We genetically manipulated the intrinsic DNA curvature and observed an altered mutation rate consistent with the genome-wide data. We showed that this effect is due to increased mutation rate, likely due to increased exposure to mutagens, and not due to differential efficacy of repair machinery. Taken together, our study extensively expands our knowledge of elements that regulate mutation rate by integrating the valuable information of DNA shape, and develops a new framework to understand evolution and tumorigenesis at a nucleotide resolution.

## METHODS

### Characterization of the mutational landscape of *URA3*

A haploid *S. cerevisiae* strain derived from the W303 background, GIL104 (*MAT**a** URA3*, *leu2*, *trp1*, *CAN1*, ade2, *his3*, *bar1Δ::ADE2*), was used to characterize the mutational landscape of *URA3.* Cells from a single colony were cultured in 5 ml SC media with uracil dropped-out (SC-uracil) at 30°C for 24 hours. Cells were then transferred into 5 ml fresh SC media (at an initial OD_660_ ∼0.1) and grown for 24 hours to accumulate mutations. ∼5.0×10^7^ cells were spread onto SC-uracil plates containing 1 g/l 5-FOA to select for loss-of-function mutants of *URA3.* A total of ∼1,000 *ura3* variants were isolated from 5-FOA plates and were Sanger sequenced separately. PCR and Sanger sequencing primers are listed in **Table S5**.

### Calculation of the mutation rate and the values of DNA properties in *URA3* and *CAN1*

We identified a total of 452 mutations in *URA3* (**Table S1**), including 5 synonymous mutations, 293 missense mutations, and 154 nonsense mutations. We focused on these 154 nonsense mutations in this study for the sake of accuracy in estimating mutation rate. To be specific, we need to count the number of potential loss-of-function mutation sites, which would be used to normalize the number of observed mutations and hence to calculate the mutation rate. The number of potential loss-of-function missense mutations was difficult to estimate because it remains elusive which missense mutations lead to a loss of function and which do not. Mutation rate was determined using overlapping windows with size equal to *L* nucleotides (*L*= 10, 20 …, or 100 bp, **Fig. S3A**). The window slid for 10 nucleotides each movement. The value of a DNA shape feature were calculated based on the frequencies of all 16 possible combinations of dinucleotide in a region, following previous studies [19, 23]. The value of each dinucleotide for each DNA shape feature was obtained from the DNA ‘PROPERTY’ database [46] and was shown in **Fig. S3B**. GC content was calculated with a Perl script.

### Estimation of nucleosome occupancy

The wild-type *S. cerevisiae* strain (BY4741 *URA3*) was grown to log-phase in YPD (1% yeast extract, 2% peptone, and 2% dextrose) liquid medium. We performed nucleus isolation, micrococcal nuclease (MNase) digestion, and chromatin preparation as described previously [47], with the following modifications. We adjusted NP-S buffer to 0.5 mM spermidine, 0.075% (v/v) NP-40, 50 mM NaCl, 50 mM Tris-HCl pH 7.5, 5 mM MgCl_2_, and 5 mM CaCl_2_, and used 100 units of MNase to digest the nuclei for 5 minutes. We performed Protease K digestion and exacted the core particle DNA. Paired-end libraries were constructed using Illumina-compatible DNA-Seq NGS library preparation kit from Gnomegen and were sequenced with Illumina HiSeq 2500 (PE125, paired-end 2 × 125 bp). ∼10.6 million clean reads were aligned to the *S. cerevisiae* genome using bowtie2 with default parameters [48]. Nucleosome occupancy of a nucleotide was defined as the number of read pairs uniquely mapped to the genome region covering the nucleotide. The raw sequencing data of MNase-seq have been deposited to the Genome Sequence Archive [49] in BIG Data Center (http://bigd.big.ac.cn/gsa), Beijing Institute of Genomics, Chinese Academy of Sciences, under accession number CRA000570.

### Generation and analyses of the *URA3* variants

We designed four synonymous variants of *URA3* with different intrinsic DNA curvature (**Tables S3-4**). We estimated the minimum free energy (MFE) for all 20 nucleotide windows in the coding sequence with RNAfold [50], and defined the average MFE of them as the strength of the RNA secondary structure of a variant. Codon adaptation index (CAI) was calculated following our previous study [51]. Four *URA3* variants were synthesized by Wuxi Qinglan Biotech and the wild-type *URA3* DNA sequence was amplified from S288C. Primers are listed in **Table S5**. Each of the five variants was introduced into the chromosomal location of *URA3* in BY4741 (*MAT***a** *his3*Δ*1 leu2*Δ*0 met15*Δ*0 ura3*Δ*0*) with homologous recombination.

We used electrophoretic mobility shift assay to confirm the difference in intrinsic DNA curvature of the five synonymous variants. We loaded an equal amount of PCR products of five variants into a 12% native polyacrylamide gel. We performed the electrophoresis experiement in the TBE buffer (89 mM Tris, 89 mM boric acid, and 2.5 mM EDTA, pH 8.0) for 12 hours at 120 V.

Total RNA was extracted with hot acidic phenol (pH < 5.0) and was reverse transcribed with the GoScript^™^ reverse transcriptase. Quantitative PCR (qPCR) was carried out on the Mx3000P qPCR System (Agilent Technologies) using Maxima SYBR Green/ROX qPCR Master Mix. *ACT1* was used as the internal control. Primers used are listed in **Table S5**.

The variance-to-mean ratio of the numbers of colonies on the plates was much greater than 1 for each variant (**Fig. S11**), indicating that the number of colonies does not follow a Poisson distribution [33]. This suggests that the observed mutations most likely occurred in the liquid culture instead of on the plates. We used the non-parametric Mann-Whitney *U* test to compare the number of colonies among these strains. We also estimated the relative mutation rates in these variants from *p*_0_, the proportion of cultures with no mutants, in the wild-type background with the following equation [33].

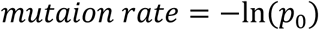

### Estimation of mutation rate in yeast mutation accumulation (MA) lines

A previous study identified ∼1,000 single nucleotide mutations by sequencing the genomes of five MA lines of a mismatch repair-deficient *S. cerevisiae* strain (BY4741 *msh2::kanMX4*) [27]. The mutation data from this study was used because the efficacy of purifying selection in MA experiments [17, 22] was further reduced in mutators. We analyzed the mutations supported by ≥ 20× coverage and retrieved 882 single nucleotide mutations that were identified in at least one of the five replicates from this study. As a control, we chose 882 random sites in the rest of the yeast genome and defined them as the pseudo-mutation sites. We calculated the average intrinsic DNA curvature around these pseudo-mutation sites and repeated this procedure for 1,000 times. *P* values were calculated as the fraction of pseudo-mutation sets exhibiting a smaller average intrinsic DNA curvature than that of the observed mutation sites among 1,000 permutations.

### Estimation of mutation rate in human cancer cells

The data of SNPs in cancer cells were retrieved from The Cancer Genome Atlas (TCGA) database [29]. Chromosomal sequences surrounding these SNPs were retrieved from Ensembl release 87 (www.ensembl.org). When multiple projects for a cancer type exist, we combined all SNPs in these projects. On average, ∼100,000 SNPs were identified in a cancer type. For each cancer type, we calculated the average intrinsic DNA curvature of the flanking DNA sequences of all SNPs (from 50 bp upstream to 50 bp downstream of each SNP). We also randomly chose the same number of sites in the human genome and calculated the average intrinsic DNA curvature of their flanking sequences similarly. This procedure was repeated 1,000 times to obtain the distribution of the expected average intrinsic DNA curvature. *P* values were calculated as the fraction of sets of random sites exhibiting a smaller average intrinsic DNA curvature than that of the observed SNP sites, among 1,000 permutations. In TCGA, different methods were used to identify mutations (Mutect, Muse, Somaticsniper, and Varscan). Our conclusion held under each kind of method used in calling SNPs (**Figs. S4-6**).

### Data retrieval

Protein-protein interaction (PPI) data in yeast were downloaded from *Saccharomyces* Genome Database [52]. Lists of essential genes and haploinsufficient genes were retrieved from a previous study [53]. Genes leading to significant growth reduction upon deletion were identified in a previous study with Bar-seq [54]. Duplicate genes in the yeast genome were defined in a previous study [55]. PPI data in humans were downloaded from Biogrid [56]. Human essential genes were retrieved from two previous studies [57, 58], respectively. The list of haploinsufficient genes in humans were retrieved from a previous study [59].

## DECLARATIONS

### Ethics approval and consent to participate

Not applicable.

### Consent for publication

Not applicable.

### Availability of data and material

The raw sequencing data have been deposited to the Genome Sequence Archive in BIG Data Center (http://bigd.big.ac.cn/gsa) under accession number CRA000570.

## Authors’ contributions

C.D., L.B.C., X.H., and W.Q. designed the experiments. C.D., Q.H., and X.C. performed the experiments. C.D., Q.H., and W.Q. analyzed the data. C.D., S.W., L.B.C., and W.Q. wrote the manuscript.

## Acknowledgements

We thank Yuliang Zhang for technical support in data analysis, and Mengyi Sun and Jian-Rong Yang for critical reading of the manuscript.

